# Isotonic medium treatment limits burn wound microbial colonization and improves tissue repair

**DOI:** 10.1101/2024.10.29.620892

**Authors:** Adam Horn, Andrew S. Wagner, Yiran Hou, Jocelyn C. Zajac, Alexandra M. Fister, Zhili Chen, Joana Pashaj, Mary Junak, Nayanna M. Mercado Soto, Angela Gibson, Anna Huttenlocher

## Abstract

Burn injuries undergo a complex healing process in which progressive spreading of epithelial damage can lead to secondary complications such as wound infection, which is a major driver of mortality among burn patients. We recently reported that burning larval zebrafish triggers dysregulated keratinocyte dynamics compared to mechanical injury. Here, we investigate keratinocyte behavior following burn injury and the subsequent potential for microbial colonization of burn wounds over time. Real-time imaging, coupled with tracking of photoconverted cells, revealed that early keratinocyte motility contributes to the spread of epithelial damage beyond the initial site of burn injury and that increased epithelial damage was associated with wound colonization by the fungal pathogen *Candida albicans*. Modulating osmotic balance by treating larval zebrafish with isotonic medium limited the spread of epithelial damage and reduced microbial colonization of burn wounds. Using cultured human skin, we found that topical treatment with isotonic solution (saline) similarly prevented the spread of epithelial damage over time. These findings indicate that keratinocyte behavior contribute to burn wound progression in larval zebrafish and link keratinocyte dynamics to microbial colonization of burn wounded tissue.

## Introduction

The epithelium provides a protective barrier that separates the external environment from internal tissues. While epithelial tissue is well-adapted to repair from mechanical damage, burn wounds often result in long-term physical and psychosocial impacts on patients. Inefficient repair of burn wounded tissue can lead to permanent scarring, chronic pain, and impaired skin function.^[1–3]^

Burn wound conversion is a characteristic feature of burn injury and describes the phenomenon of superficial burns converting to deeper burns over time.^[4]^ This concept originates from the description by Jackson of the zones of burn injury.^[5]^ In this framework, the central region of irreversibly-damaged necrotic tissue is termed the zone of coagulation. This region is surrounded by a zone of threatened, but viable, damaged tissue referred to as the zone of stasis. A recoverable zone of hyperemia exists at the periphery of the wound. Burn wound conversion describes the process by which the zone of stasis converts from viable to irreversibly-damaged, nonviable tissue, thereby expanding the zone of coagulation in both surface area and depth.^[4, 6]^ Limiting this progressive spreading of epithelial damage thereby prevents the death of otherwise viable tissue in the zone of stasis and increases the spontaneous healing potential of burn wounds. Although therapeutic avenues to limit the spread of burn damage have been explored, there remains a need to understand the underlying mechanisms that promote burn wound conversion.

While spread of epithelial damage is detrimental to burn wound healing, burn wound infection is a leading cause of patient mortality.^[7–9]^ The prevalence of fungal infection of burn patients is rising^[10]^, and *Candida* species are among the most common fungal pathogens found colonizing burn wounded tissue.^[11, 12]^ In 2022, the World Health Organization identified *Candida albicans* as a critical priority pathogen, highlighting the growing threat of fungal infections worldwide, especially in resource poor regions where fungal infection of burn wounds may remain undiagnosed.^[13]^ Importantly, burn patients are susceptible to delayed infection, which can occur days or weeks after the burn injury itself. This is in part due to long-lasting epithelial barrier disruption and aberrant antimicrobial activity in burned tissue, which itself is a primary site of microbial colonization.^[14]^ This apparent link between the epithelial response to burn injury and wound infection led us to hypothesize that early keratinocyte behavior contributes to burn wound conversion and enables delayed microbial colonization of wounded tissue.

To investigate this hypothesis, we used the larval zebrafish model. Larval zebrafish have been widely adopted to investigate the mechanisms of tissue repair and regeneration.^[15]^ Unlike common mammalian models used to investigate burn wound healing, larval zebrafish are optically transparent. This feature enables whole organism intravital imaging throughout the duration of wound healing allowing for investigation of the tissue response to burn injury at the cellular level. In the larval stage, zebrafish have a stratified epithelium composed of two layers of keratinocytes and a collagen rich dermis^[16]^, providing an anatomically simplified model to investigate cellular behavior in response to burn injury. We previously developed a method of burning larval zebrafish and showed that zebrafish recapitulate key features of burn wound pathology in humans including chronic inflammation, dysregulated extracellular matrix remodeling, and impaired sensory neuron regeneration.^[17–19]^ Further, we identified excessive keratinocyte motility as a driver of burn wound pathology.^[19]^

Taking advantage of the unique benefits of the larval zebrafish, we sought to investigate how the early epithelial response to burn injury contributes to pathology, including microbial colonization of wounded tissue. We found that dysregulated keratinocyte migration leads to the spread of burn wound damage over time, increasing cell death in a region of tissue analogous to the zone of stasis. This long-term disruption of epithelial barrier function was associated with increased colonization of the fungal pathogen *C. albicans*. Preventing keratinocyte motility by treating burned larvae with medium isotonic to their interstitium limited the spread of tissue damage and reduced burn wound colonization by *C. albicans*. Finally, we applied our finding that isotonic medium treatment limits the spread of burn damage in zebrafish to cultured human skin. Topical treatment of human skin with isotonic solution (saline) prevented an increase in burn depth over time suggesting that further study of early saline treatment may provide a therapeutic avenue to improve burn wound healing potential in patients.

## Materials and Methods

### Animal Use Ethics

Zebrafish studies were carried out in accordance with the recommendations from the Guide for the Care and Use of Laboratory Animals. All zebrafish experiments performed were approved by the University of Wisconsin – Madison Research Animals Resource Center under the Protocol M005405-R02.

### Zebrafish handling and maintenance

Adult and larval zebrafish were maintained as previously described^[20]^. Briefly, adult fish were maintained on a 14 hours light – 10 hours dark cycle. Following breeding, embryos were transferred to E3 medium (5 mM NaCl, 0.17 mM KCl, 0.44 mM CaCl_2_, 0.33 mM MgSO_4_, 0.025 mM NaOH, and 0.0003% methylene blue) and maintained at 28.5°C until 3 days post-fertilization. When necessary, larvae were screened for the expression of the appropriate transgene indicated by positive fluorescence using a Zeiss Zoomscope EMS3/SyCoP3 with a Plan-NeoFluar Z objective. For example, expression of the nuclear localized Dendra transgene was confirmed prior to experimentation by examining larvae for the presence of green fluorescence. For all experiments, 3 days post-fertilization larval zebrafish were anesthetized in 0.2 mg/mL tricaine (ethyl 3-aminobnzoate) in E3 medium prior to use. Following burn injury, larvae were returned to 28.5°C until imaging.

### Burn injury of larval zebrafish

To perform burn injury, zebrafish larvae were transferred to a 60 mm tissue-culture treated dish containing 0.2 mg/mL tricaine in E3 medium. Unless it is indicated that larvae were infected following injury, all burn wounds were performed in sterile medium to prevent wound contamination. A fine tip cautery pen (Geiger Instruments, Delasco Item #250) with angled loop (Geiger Instruments, Delasco – Item #201C) was used to burn the caudal fin of larval zebrafish. Burns were performed until the wounded region reached halfway to the posterior notochord boundary to avoid damage to the notochord. For multi-day experiments, larvae were washed into fresh medium immediately following burn injury and maintained at 28.5°C until imaging. Isotonic medium was prepared by supplementing control E3 medium with NaCl (Sigma-Aldrich) to a final solute concentration of 270 mOsm. For experiments involving isotonic medium treatment, larvae were only exposed to isotonic medium at the time of burn injury. Unless otherwise indicated, treatment with isotonic medium was carried out for 6 hours post-burn, at which time larvae were washed into fresh E3 medium. For cases in which imaging was performed fewer than 6 hours post-burn, treated larvae were imaged in isotonic medium.

### Photoconversion of larval zebrafish

Photoconversion of *Tg(Krt4-H2B-Dendra2)* larvae was performed using an Olympus FluoView FV-1000 laser-scanning confocal microscope equipped with a NA 0.75/20X air objective. The stable conversion of native green *Dendra2* fluorescence to red fluorescence was carried out using a 405 nm laser set to 10 µs/pixel dwell time and 45 second total stimulation time. Keratinocyte photoconversion was limited to a specific region of interest, depending on the given experiment. For all experiments, images were taken immediately prior to and after photoconversion to ensure a successful shift in Dendra2 fluorescence. To define the region used for photoconversion, a region of interest representing zone 1 or zone 2 was manually drawn. Photoconversion of anesthetized larvae was performed immediately following burn injury in all cases, with larvae being washed with either control E3 or isotonic E3 medium and returned to 28.5°C until imaging. Photoconversion data were quantified by measuring the thresholded area of the red (photoconverted) channel and normalizing relative to the photoconverted area present at 0 hours post-burn.

### Immunofluorescence and quantification of caspase staining

Larval zebrafish fixation and staining was adapted from Santos et al. 2018.^[21]^ Larvae were fixed using 4% paraformaldehyde in PBS. The following antibodies were used: 1° Rb – α-active caspase 3 (BD Pharmingen, Cat: 559565, 1:500); 2° Dk – α-Rb-IgG Dylight 550 (Invitrogen, SA5-10039, 1:300). Active-caspase area was quantified by manually thresholding for positive-fluorescence.

### Preparation of *Candida albicans* and zebrafish infection

*C. albicans* strains used in this study can be found in Table 1 below. *C. albicans* were activated on YPD agar plates (1% yeast extract, 2% peptone, 2% dextrose, 2% agar) from glycerol stocks. One day prior to infection, overnight cultures in liquid YPD media were started and left to incubate for 16 hours while shaking at 225 rpm at 30°C. For infection experiments, zebrafish larvae were exposed to *C. albicans* diluted in E3 medium without methylene blue to a final concentration of 1x10^7^ cells/mL beginning immediately after burn wounding. Larvae were incubated with *C. albicans* on an orbital shaker for 1 hour following burn injury before being washed into fresh E3 medium. To assess colony forming units, fungal stocks prepared in either control or isotonic medium were serially diluted and spread on YPD agar plates. Plates were incubated for 24 hours at 30°C before colony forming units were counted and used to determine viability.

**Table 1:** Fluorescent-tagged *Candida albicans* strains.

| Strain | Parent | Genotype | Source |
| --- | --- | --- | --- |
| Caf2-FR | Caf2.1 | $\Delta ura3::imm434/URA3 P_{ENO1}-iRFP-NATR$ | [22] |
| Caf2-dTomato | Caf2.1 | $\Delta ura3::imm434/URA3 P_{ENO1}-dTomato-NATR$ | [23] |
| SC5314-mNeon | SC5314 | $P_{ENO1}-NEON-NAT$ (Wild Type) | [24] |

### Quantification of immune cell recruitment

In all cases, neutrophils and macrophages were identified by the presence of a fluorescence transgene driven by a cell type-specific promoter. For experiments involving exposure to *C. albicans*, *Tg(LyzC-BFPxMpeg1.1-mCherry)* larvae were used. Quantification of neutrophil and macrophage recruitment at each time point was performed by measuring the total thresholded area of fluorescently labeled cells, respectively. For quantification of immune cell recruitment following sterile burn injury, *Tg(LyzCH2B-mCherryxMpeg1.1H2B-GFP)* larvae were used and neutrophil or macrophage nuclei were manually counted using the multi-point tool in FIJI 2.9.0. Only nuclei in the wound region, defined as caudal fin tissue distal the posterior notochord boundary, were quantified.

### Burn injury of cultured human skin

De-identified human skin was obtained from patients undergoing elective plastic surgery when the skin would otherwise have been discarded at our institution. The de-identified samples were exempt from the regulation of University of Wisconsin-Madison Human Subjects Committee Institutional Review Board, and no patient data was collected. Skin was processed and burned within an hour after surgery, as previously described^[25]^. Briefly, subcutaneous fat was removed from the tissue, leaving the dermis intact, and uniform rectangles of 4.2x1.5 cm were cut from the skin. A custom fabricated device was used to burn three 8 mm partial thickness contact burns per skin section by preheating the device to 150°C for 6 seconds^[25]^. Skin samples were cultured on custom elevated metal mesh inserts, allowing skin to be cultured at an air-liquid interface. Culture media, composed of DMEM (Gibco Laboratories, Gaithersburg, MD), 10% FBS (Gemini Bio, Sacramento, CA), 0.625 µg/ml amphotericin B (Gemini Bio, Sacramento, CA), and 100 µg/ml penicillin-streptomycin (Gibco Laboratories, Gaithersburg, MD) was changed daily and samples were kept in an incubator at 37°C and 5% CO_2_ for the duration of each experiment. Treatments were applied to burned skin using rubber O-rings adhered to skin with petroleum jelly. Liquid treatments (sterile water and saline) were both kept at room temperature prior to topical application to ensure that any change in burn wound temperature would be consistent across treatment groups. At the appropriate time point for collection, full thickness biopsies were taken by 12 mm biopsy punch with the area of burned tissue in the center and surrounding region of unburned skin being included. Samples were cryopreserved with Tissue-Tek compound (Sakura Finetek USA, Torrance). Frozen samples were cryo-sectioned to 6 µm slides and stained for lactate dehydrogenase (LDH) following the protocol developed by Gibson and Shatadal^[26]^. Sections were imaged to determine the depth of dermal injury defined by the vertical distance from the center of the burn between the cutaneous basement membrane to the lower depth of LDH staining in the dermis.

### Microscopy and image analysis

For experiments including photoconverted larvae and larvae exposed to *C. albicans*, imaging was performed using a spinning disc microscope (CSU-X, Yokogawa) with a confocal scanhead on a Zeiss Observer Z.1 inverted microscope, either a Plan-Apochromat NA 0.8/20X objective or a Plan-Apochromat NA 1.4/63X objective, and a Photometrics Evolve EMCCD camera. ZEN 2.6 software was used for acquisition. Larvae exposed to far-red expressing *C. albicans* were imaged using a Nikon Eclipse TE300 inverted fluorescent microscope with a Plan Apo NA 0.45/10X objective. Image acquisition was performed using Nikon NIS-Elements software. Time-series, bright field imaging of burned larvae was performed on a Zeiss Zoomscope EMS3/SyCoP3 with a Plan-NeoFluar Z objective running Zen 3.7. In all cases, anesthetized larvae were mounted in 2% low-melting point agarose on a 35 mm glass-bottom dish (CellVis). All image processing and analysis was performed using FIJI 2.9.0. Unless otherwise indicated fluorescent images are presented as maximum intensity z-projections. This method of projecting 3D data to a 2D image works by identifying the brightest voxel (with maximum fluorescence intensity) at a given XY position through the entire depth, in z, of an image stack. The brightest voxel at each XY position is used to generate the 2D projection regardless of its original position in z-depth. Methods used for quantification of imaging data are detailed within the appropriate section above.

### Statistical analysis

All statistical analysis and graphing was performed using GraphPad Prism version 9.0.0. The D’Agostino and Pearson normality test was used to determine the appropriate statistical test based on normal distribution of data. For comparison of two means, independent t-tests were used for normally distributed data, whereas Mann-Whitney tests were used for nonparametric data. For multiple comparisons, one-way ANOVA with Tukey’s multiple comparison test was used for normally distributed data with Kruskal-Wallis with Dunn’s multiple comparison test used for nonparametric data. Two-way ANOVA with Sidak’s multiple comparison test was used when appropriate depending upon the number of independent variables in each experiment. Unless otherwise indicated, all data are represented as mean ± SEM (standard error) with p<0.05, indicated by asterisk, used to determine statistical significance. All experiments were performed with at least 3 independent biological replicates defined as either separate breeding clutches of zebrafish or human subjects, with the N-value used for statistical analysis indicated within the relevant figure legend. When possible, measures such as blinded analysis were performed to limit potential biases.

## Results

### Burn injury causes local and long-lasting epithelial damage in larval zebrafish

To assess the epithelial response to injury, we burned 3 days post-fertilization larval zebrafish using a hand-held cauterizer. This method allows for reproducible, localized epithelial damage, which is evident by the entry of cell impermeable dye only in tissue directly impacted by burn injury (**Figure 1A**). Immediately following burn injury, the presence of localized damage is visible by a change in tissue opacity (**Figure 1B**, blue outline). Epithelial damage continues to accumulate in the burned region until 6 hours post-burn (hpb) (**Figure 1B**, white arrow). By 24 hpb, the remodeling phase of tissue repair has begun, with visible epithelial damage being cleared and fin regrowth beginning. Full fin regrowth, indicating successful wound healing occurs at roughly 72 hpb (**Figure 1B**). We recently reported that excessive keratinocyte motility occurs within the first hour following burn injury, and motility can be prevented when larvae are burned in the presence of medium isotonic to the interstitium (**Figure 1C**).^[19]^ This is thought to occur by altering the osmotic balance. In homeostatic conditions, the surrounding hypotonic freshwater environment results in fluid influx upon epithelial barrier disruption. Subsequent keratinocyte swelling is a pro-migratory signal.^[27, 28]^ In contrast, an isotonic environment prevents fluid influx and cell swelling thereby inhibiting keratinocyte migration.^[27]^ Treatment of larval zebrafish with isotonic medium prior to burn injury improved gross fin morphology, suggesting a reduction in overall epithelial damage (**Figure 1D**). From this, we hypothesized that aberrant keratinocyte behavior may contribute to the spread of epithelial damage over time, and that isotonic medium treatment may provide a means to prevent long-term disruption to epithelial barrier function.

**Figure 1:**
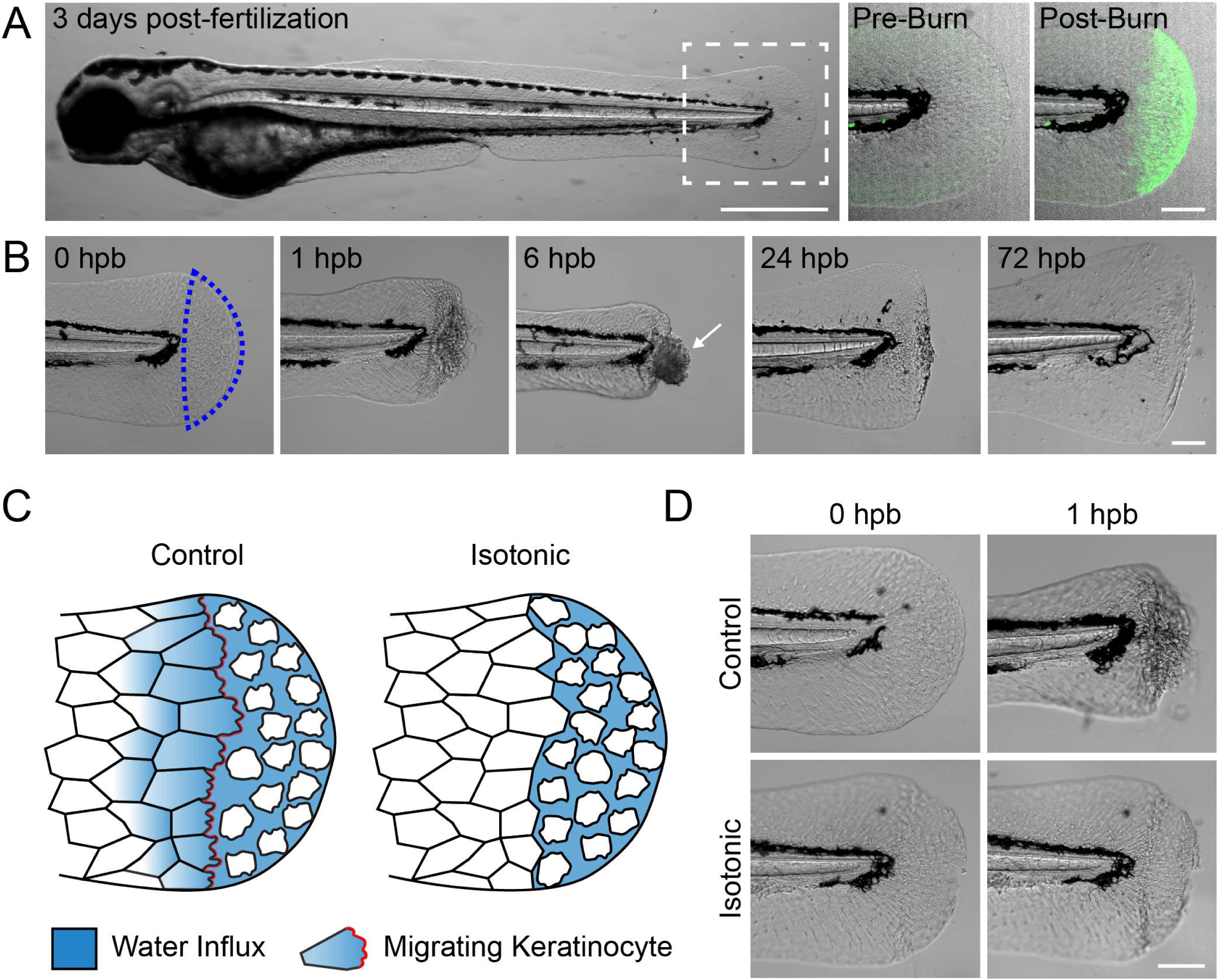
Burn injury generates local epithelial damage in larval zebrafish. (A) Larval zebrafish 3 days post-fertilization. Dashed box indicates the tailfin region used for burn experiments. Scale bar = 500 µm. At right, representative images show a tailfin pre-and post-burn in the presence of cell impermeable FM dye. Increased dye fluorescence post-burn indicates localized tissue damage. Scale bar = 100 µm. (B) Time-series showing a larval zebrafish tailfin at the indicated time, hours post-burn (hpb). Scale bar = 100 µm. (C) Schematic illustrating the effect of isotonic medium on keratinocyte dynamics after burn injury. In control (hypotonic) conditions, net fluid influx into damaged tissue and subsequent swelling of wound-edge keratinocytes stimulates a migratory response. Lack of fluid influx in isotonic conditions prevents keratinocyte swelling and cell migration. (D) Images showing control and isotonic medium treated larvae at the indicated time post-burn. Scale bar = 100 µm.

### Keratinocyte motility contributes to burn wound conversion

To determine the fate of epithelial keratinocytes following burn injury we used *Tg(Krt4-H2B-Dendra2)* zebrafish, which express nuclear-localized Dendra2 under a pan-keratinocyte promoter. Dendra2 is a photoconvertible protein, allowing for real-time switching of fluorescence from green to red upon exposure to 405 nm light (**Figure 2A**).^[29]^ Through this process, it is possible to track the fate of keratinocytes from a specific region of tissue over time. In human burn injury, the zone of coagulation is composed of necrotic tissue. Thus, we first decided to photoconvert only cells in the region of tissue immediately affected by burn, referred to here as zone 1 to determine their fate as burn injury progresses (**Figure 2A, B**). Following injury, photoconverted nuclei are rapidly lost from burned tissue with only 15 ± 2.5% of cells remaining by 6 hpb (**Figure 2C**), indicating widespread keratinocyte death over time. Similar to larvae burned in control E3 medium, we found that only 26 ± 4.9% of cells in zone 1 remained at 6 hpb following treatment with isotonic medium (**Figure 2B, C**). To confirm death of cells in zone 1, we examined this region 24 hpb and found nearly zero remaining keratinocytes regardless of treatment (**Supplemental Figure 1A, B**). Following this, we photoconverted cells in the region of tissue directly adjacent to that affected by burn, which is analogous to the zone of stasis and referred to here as zone 2 (**Figure 2A**). Similar to zone 1 photoconversion, when zone 2 keratinocytes from control larvae were tracked, we noted a loss of cells over time, albeit with different kinetics (**Figure 2D**). While approximately 50% of zone 2 keratinocytes were lost 24 hpb in control larvae, this stabilized such that 38 ± 6.2% of the original keratinocytes from zone 2 remained 72 hpb (**Figure 2E**). This finding demonstrates that progressive spreading of epithelial damage, a component of burn wound conversion, is a feature of the epithelial response to burn injury in larval zebrafish. To determine whether treatment with isotonic medium limits burn wound conversion, we photoconverted zone 2 tissue in larvae exposed to isotonic solution (**Figure 2D**). Strikingly, we found that 93 ± 5.8% of zone 2 keratinocytes remained viable 72 hpb demonstrating that cells in this zone were protected from death (**Figure 2E**). While necrotic death is observed in the zone of coagulation, and likely explains the rapid loss of photoconverted keratinocytes we observe in zone 1, apoptosis is thought to contribute to cell death during burn wound conversion in the zone of stasis.^[30, 31]^ Therefore, we next probed for apoptotic death using antibodies against active-caspase 3 in order to determine whether isotonic medium protects cells from this mode of death. However, we found no difference in the amount of apoptotic cells between control and isotonic treated fish (**Supplemental Figure 1C, D**).

**Figure 2:**
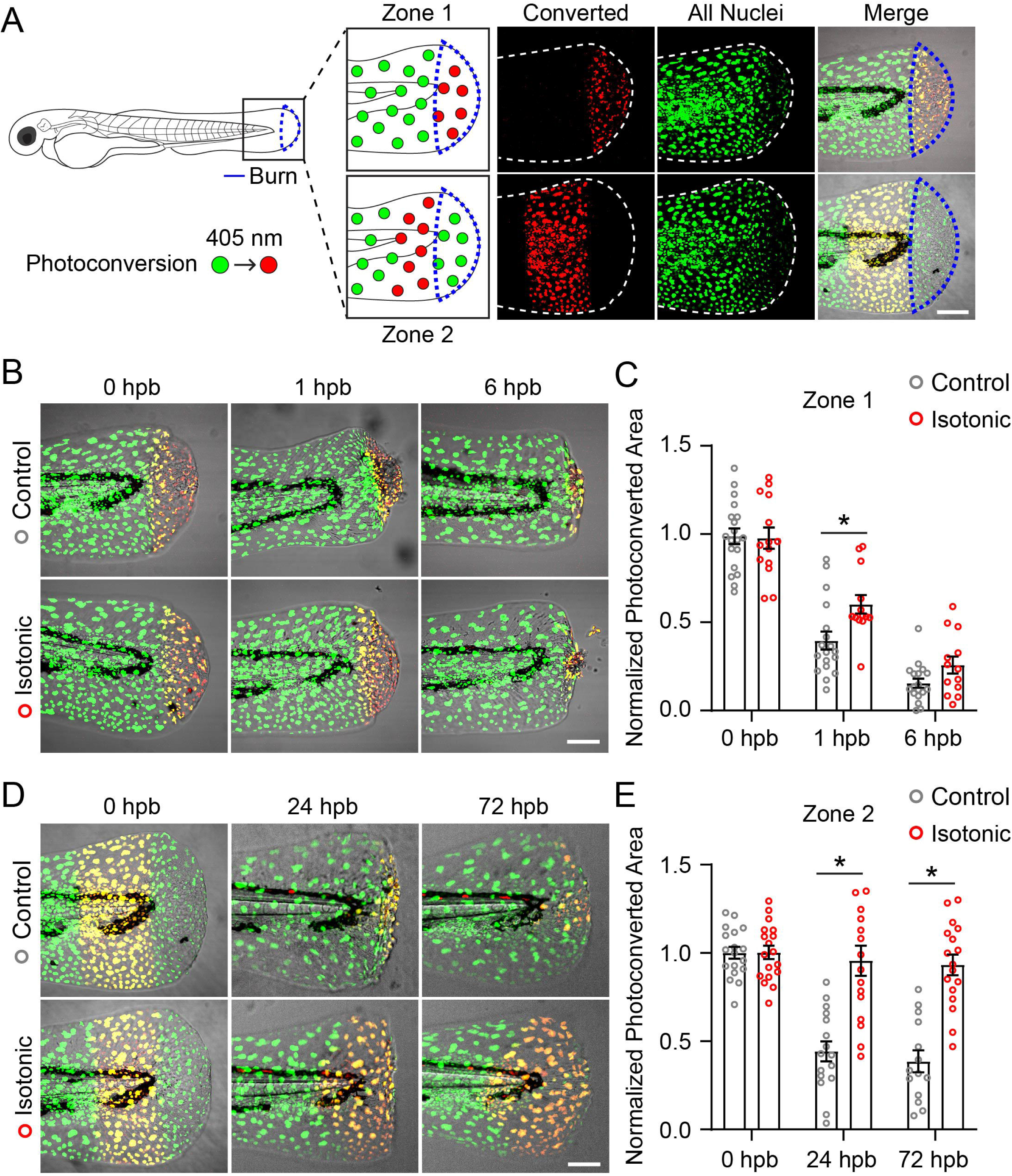
Keratinocyte motility contributes to spread of epithelial damage after burn. (A) Schematic of photoconversion experiment. Keratinocyte nuclei expressing Dendra2 (green) can be photoconverted (red) by exposure to 405 nm light. Photoconversion was performed in the indicated region immediately following burn injury indicated by the blue outline. In merged images, photoconverted nuclei appear yellow due to residual green Dendra2 fluorescence. (B) Images showing control and isotonic medium treated larvae after photoconversion of zone 1 tissue. (C) Quantification of zone 1 photoconverted area at the indicated time, hours post-burn (hpb). Data are normalized to photoconverted area 0 hpb. N ≥ 18 control and N ≥ 13 isotonic treated larvae. (D) Images showing control and isotonic medium treated larvae after photoconversion of zone 2 tissue. (E) Quantification of zone 2 photoconverted area at the indicated time, hours post-burn (hpb). Data are normalized to photoconverted area 0 hpb. N ≥ 14 control and N ≥ 16 isotonic treated larvae. Scale bars = 100 µm. Asterisk indicates p<0.05 by two-way ANOVA.

### Treatment with isotonic medium limits *Candida albicans* colonization of burned tissue

Due to the role of keratinocytes in maintaining epithelial barrier integrity and our finding that keratinocyte motility contributes to burn wound conversion, we next investigated what impact keratinocyte behavior has on microbial colonization of burn wounded tissue. To do this, we developed a method for exposing burned larval zebrafish to the fungal pathogen *Candida albicans*. Immediately following burn injury, larvae were bathed in mNeon-expressing *C. albicans* for 1 hour with wound colonization quantified 24 hpb. Wound colonization was distinguished based on the presence of *C. albicans* at the interface between viable and non-viable tissue^[10, 32]^. Comparing burned to unwounded larvae following *C. albicans* exposure revealed that *C. albicans* was unable to colonize epithelial tissue in the absence of burn injury and that successful colonization of wounded epithelium was specifically localized to the site of tissue damage (**Figure 3A**). While some *C. albicans* was identified near the developing gills in both burned and unwounded fish, this was minimal and was entirely cleared in both populations within a day of exposure. Importantly, *C. albicans* was viable within colonized tissue and able to develop hyphae, which is essential for potential tissue invasion (**Supplemental Figure 2A**).^[33]^

**Figure 3:**
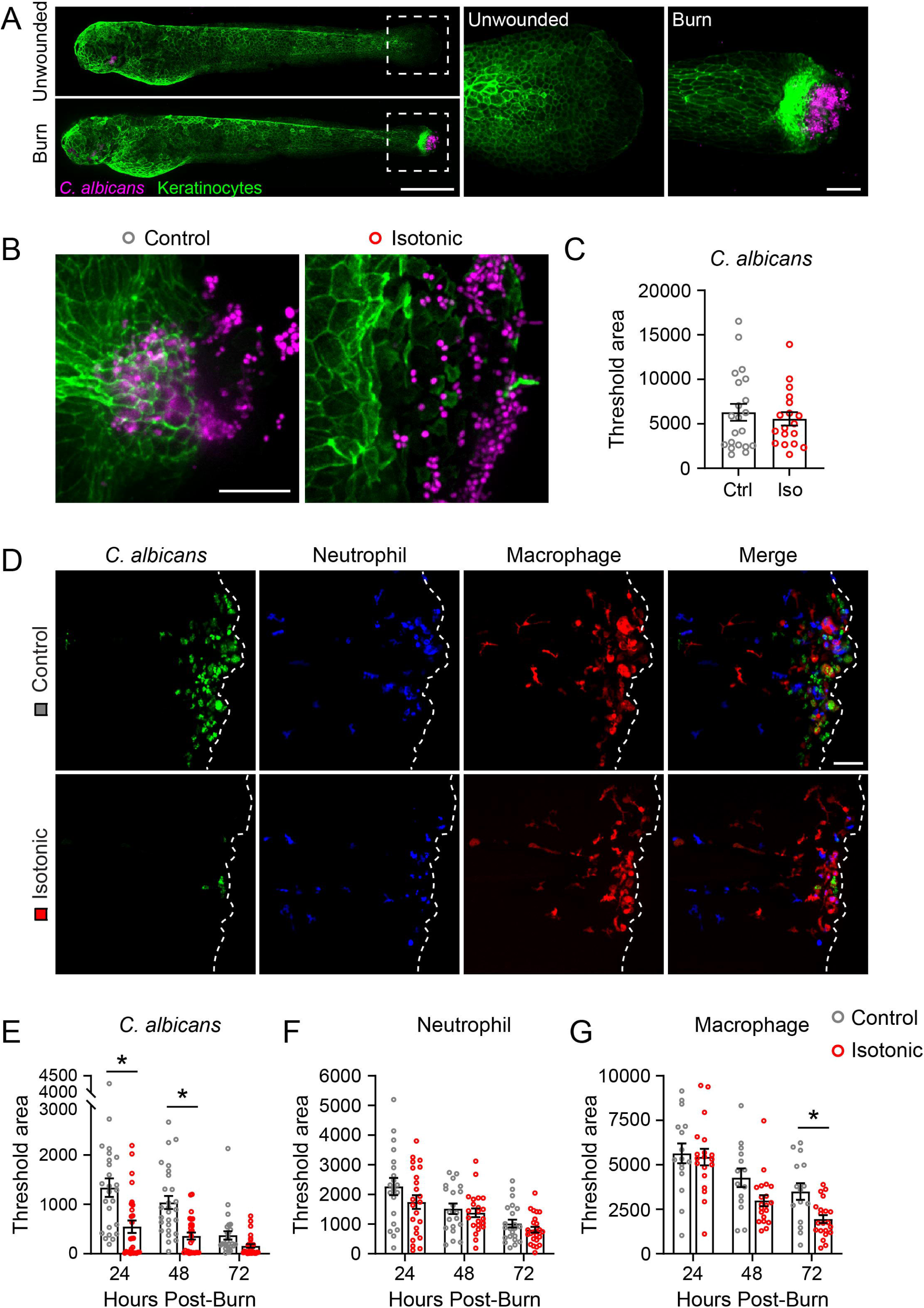
Isotonic medium treatment reduces microbial colonization of burned tissue. (A) Unwounded or burn wounded *Tg(Krt4-UtrCH-GFP)* larval zebrafish, labeling epithelial keratinocytes, after 1 hour exposure to RFP-tagged *C. albicans*. Dashed box shows area of inset to the right. Scale bar = 500 µm (whole larvae) and 50 µm (inset). (B) *C. albicans* colonization of control and isotonic medium treated tissue 1 hpb. (C) Quantification of *C. albicans* colonization 1 hpb. N = 21 control and 18 isotonic treated larvae per condition. (D) mNeon-tagged *C. albicans* colonization of *Tg(LyzC-BFPxMpeg1.1-mCherry)* larvae 24 hpb. Dashed line indicates tailfin boundary. (E-G) Quantification of larvae burned as in D showing *C. albicans* colonization (E, N≥26 control and 27 isotonic treated larvae), neutrophil recruitment (F, N≥20 control and 22 isotonic treated larvae), and macrophage recruitment (G, N≥14 control and 19 isotonic treated larvae) over time. Scale bar = 50 µm in B, D. Asterisk indicates p<0.05 by two-tailed Mann-Whitney test (C) and two-way ANOVA (E-G).

We next sought to test whether isotonic medium alters *C. albicans* colonization. To do this, we first confirmed that treatment with isotonic medium did not affect *C. albicans* independent of the host response. We found that exposure to isotonic medium did not affect fungal viability (**Supplemental Figure 2B**) or the initial fungal burden in burn wounded tissue immediately after exposure (**Figure 3B, C**). We next burned *Tg(LyzC-BFPxMpeg1.1-mCherry)* zebrafish larvae, in which neutrophils and macrophages are fluorescently labeled, in order to get a full picture of the host response to microbial colonization. We found that larvae burned in control medium were effective in clearing *C. albicans* from the wound region by 72 hpb, with neutrophils and macrophages resolving after an initial peak at 24 hpb (**Figure 3D-G**). This suggests that an effective immune response is capable of clearing *C. albicans* before infection can occur; however, patients with severe burns are immunocompromised as a result of burn damage and are at greater risk of infection.^[34–36]^ Therefore, we asked whether targeting keratinocyte motility by treating larvae with isotonic medium affected microbial colonization, as a means to reduce the risk of infection altogether. We found that isotonic medium treatment resulted in larvae having approximately 40% of the total amount of *C. albicans* present in burn wounded tissue 24 hpb relative to controls. This decrease was maintained over the course of the healing response, with treated larvae still having significantly less *C. albicans* present even 72 hpb (**Figure 3D, E**). The decrease in fungal burden did not seem to be dependent on the host immune response evident by similar levels of neutrophils and macrophages in wounded tissue over time when compared to control larvae (**Figure 3F, G**). However, similar analysis of immune cell recruitment in the absence of *C. albicans* (sterile burn) revealed a clear difference in neutrophil recruitment, with control larvae having an average of 28.5 ± 2.0 neutrophils at the time of maximum tissue damage (6 hpb) compared with only 8.7 ± 1.6 in isotonic treated larvae (**Supplemental Figure 2C, D**). Analysis of macrophage recruitment to sterile burn injury revealed a significant difference during the period of wound remodeling, 24 hpb, with control larvae having 50.0 ± 2.5 macrophages compared to only 21.1 ± 2.1 following isotonic treatment (**Supplemental Figure 2C, E**). This opens the possibility that a dampened immune response due to isotonic treatment could aid in preventing inflammatory tissue damage.

### Early keratinocyte motility mediates *C. albicans* colonization following burn injury

While we observed that treatment with isotonic medium did not alter the initial fungal burden after one hour of *C. albicans* exposure (**Figure 3B, C**), we noted that the spatial distribution of *C. albicans* was markedly distinct. In control larvae, but not those treated with isotonic medium, *C. albicans* appeared to overlap with viable tissue corresponding to zone 2. This led us to hypothesize that isotonic medium limits fungal colonization at the site of damage by reducing burn wound conversion, maintaining epithelial barrier integrity, and thereby preventing *C. albicans* from accessing viable tissue in the zone of stasis. To test this hypothesis, we repeated our previous photoconversion experiment using *Tg(Krt4-H2B-Dendra2)* larvae, but with the addition of *C. albicans* exposure. We visualized *C. albicans* colonization 1 hpb following zone 2 photoconversion as this is the time at which keratinocyte migration has stopped. Examining control larvae revealed *C. albicans* was already present in zone 2 indicating the ability of *Candida* to colonize viable tissue (**Figure 4A**). Conversely, *C. albicans* did not spatially overlap with zone 2 keratinocytes when larvae were treated with isotonic medium (**Figure 4A**). This demonstrates that inhibiting keratinocyte motility by treatment with isotonic medium prevents colonization of viable zone 2 tissue, instead limiting *C. albicans* to colonize nonviable tissue in zone 1.

**Figure 4:**
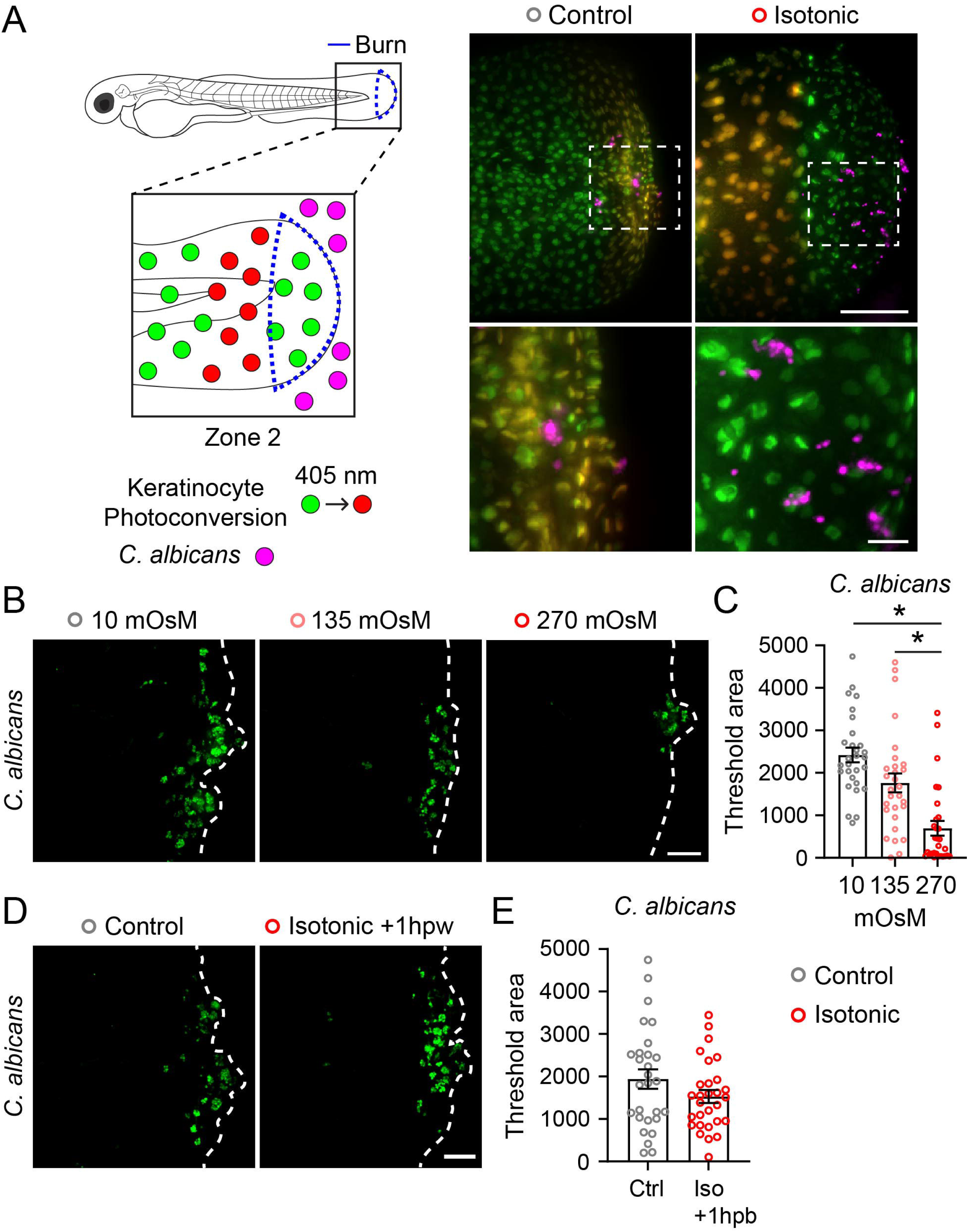
Keratinocyte dynamics enable microbial colonization of burn wounded tissue. (A) Schematic of photoconversion experiment. *Tg(Krt4-H2B-Dendra2)* larvae were photoconverted to track zone 2 keratinocytes following exposure to far red-expressing *C. albicans*. Images show spatial localization of *C. albicans* (magenta) relative to zone 2 keratinocytes (yellow) with the indicated treatment 1 hpb. Dashed box indicates region of inset shown below. Scale bar = 100 µm and 20 µm (inset). (B) *C. albicans* colonization of wound area 24 hpb. Larvae were treated with medium supplemented with NaCl to a final concentration of 10 mOsm (Control), 135 mOsm, or 270 mOsm (Isotonic). Dashed line indicates tailfin boundary. (C) Quantification of *C. albicans* colonization from B. N = 29 larvae for each condition. (D) *C.* albicans colonization of wound area 24 hpb. All larvae were burned in control medium, and treatment with isotonic medium began 1 hpb. Dashed line indicates tailfin boundary. (E) Quantification of *C. albicans* colonization from D. N = 28 control and 29 isotonic treated larvae. Scale bars = 50 µm unless otherwise indicated. Asterisk indicates p<0.05 by Kruskal-Wallis test (C) and independent t-test (E).

To directly test whether osmotic regulation of keratinocyte migration contributes to microbial colonization of viable tissue, we treated larvae with medium at an intermediate tonicity between control and isotonic medium. It has been shown that wound-induced cell migration is sensitive to extracellular tonicity in a dose-dependent manner.^[37]^ In control conditions, water used to maintain zebrafish is kept at approximately 10 mOsm while medium isotonic to zebrafish interstitial tissue is 270 mOsm. Therefore, we supplemented normal E3 medium with NaCl to an osmolarity of 135 mOsm in order to reduce, but not eliminate, keratinocyte motility. Following burn injury, we observed that larvae treated with 135 mOsm medium had an intermediate level of *C. albicans* colonization relative to control and isotonic medium treated fish at 24 hpb, supporting a role for keratinocyte migration in enabling microbial colonization (**Figure 4B, C**). In all previous experiments to this point, larvae exposed to isotonic medium were treated for 6 hours following burn, but keratinocyte migration only occurs for 1 hour. Therefore, it remained possible that the reduced colonization of viable zone 2 tissue that we observe in isotonic conditions was due to an effect of isotonic medium that is independent of keratinocyte motility. To determine if this was the case, we delayed isotonic medium treatment until 1 hpb. Assessing *C. albicans* colonization 24 hpb revealed that delayed isotonic medium treatment resulted in no difference in fungal burden relative to control fish (**Figure 4D, E**). This demonstrates the importance of early keratinocyte migration specifically in promoting increased *C. albicans* colonization of burn wounded tissue.

### Topical saline treatment improves the healing potential of burned human skin

Standard of care for initial large burn wound treatment includes administering intravenous fluids to resuscitate the patient, allowing for perfusion of damaged tissue.^[38]^ Because of the importance of fluid balance in healing burn injuries, in combination with our findings from larval zebrafish, we hypothesized that topical application of isotonic medium (saline) to human skin may improve burn wound healing.

We devised a method of testing this hypothesis by using cultured human skin, which was burned and cultured as previously described.^[25]^ Untreated skin biopsies were left exposed only to air, while treated biopsies were topically exposed to either sterile water (hypotonic solution) or saline (isotonic solution) for 1-day post-burn (1 dpb) (**Figure 5A**). Skin biopsies were taken up to 6 dpb as this coincides with the phase of burn wound expansion, and cell viability indicated by LDH staining was used to quantify burn depth (**Figure 5B, C**).^[26]^ We found that burn depth was unchanged 1 dpb regardless of treatment, highlighting that our burn wound paradigm was able to produce wounds of consistent depth. In untreated samples, we noted the expansion of burn depth over time such that by 2 dpb and extending to 6 dpb burn depth had increased from an average depth of 828 ± 50.7 µm to 1069 ± 63.6 µm and 1316 ± 94.8 µm, respectively. This represents a nearly 60% increase in burn depth over 6 days and is indicative of burn wound conversion (**Figure 5D**). Treatment of skin samples with sterile water showed reduced burn wound conversion with burn depth only increasing from 884 ± 73.4 µm to 1034 ± 68.0 µm over the first 6 days post-burn. However, treatment with isotonic saline showed no evidence of wound expansion that would indicate burn wound conversion and instead exhibited a slight decrease in wound depth. Burn depth of saline treated samples decreased from 710 ± 34.1 µm to 689.4 µm over 6 days, indicating potential wound healing (**Figure 5D**).

**Figure 5:**
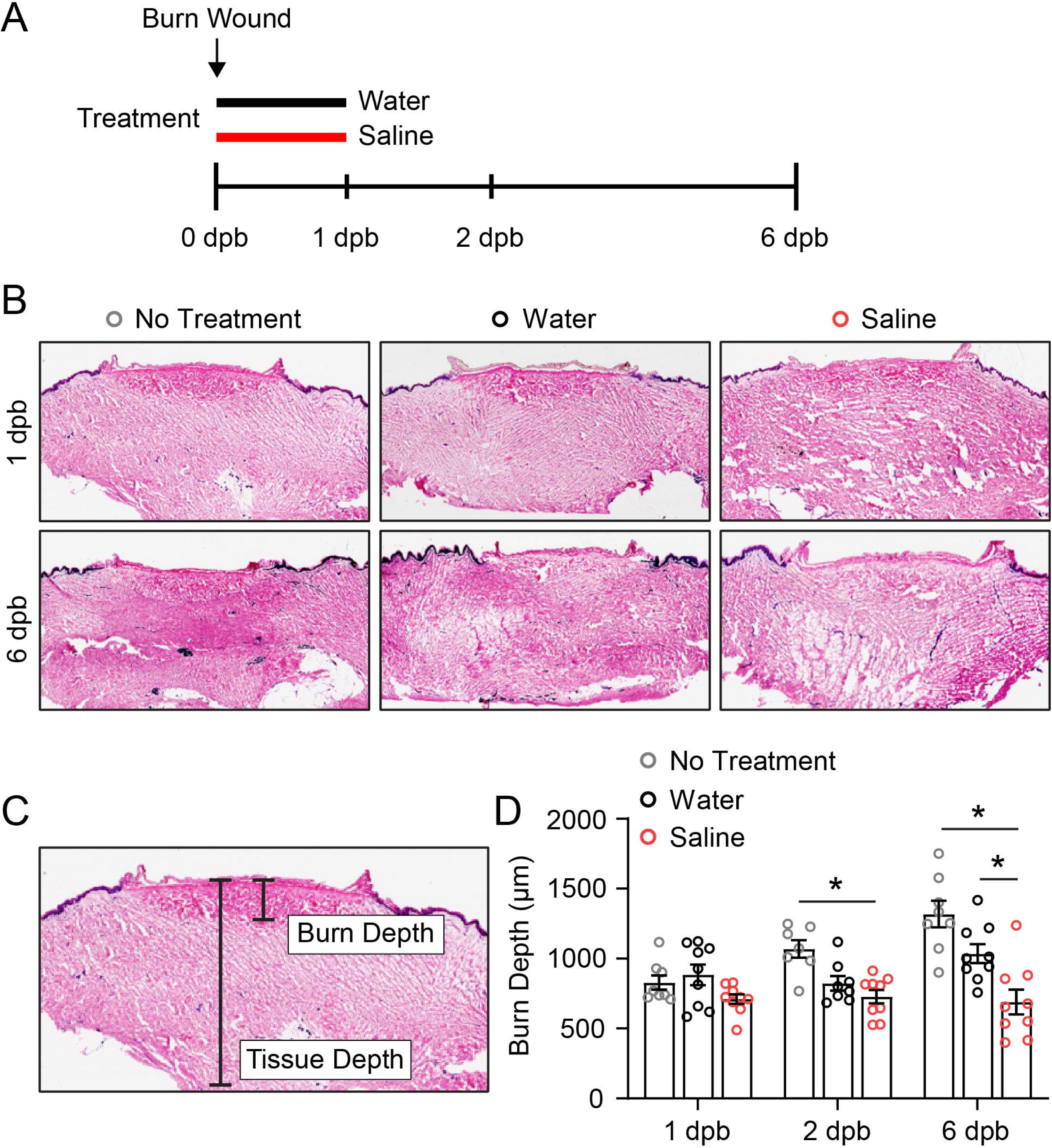
Treatment with topical saline improves burn wound healing potential of cultured human skin. (A) Experimental timeline for burn wounding of cultured human skin. The indicated treatments were applied topically for 1 day post-burn (dpb) and frozen skin sections were taken at each time point indicated. (B) Representative images of frozen skin sections stained with LDH to assess tissue viability. (C) Image showing method of quantification for burn depth. Blinded analysis was performed to measure LDH stained tissue. (D) Quantification of burn depth over time with the indicated treatment. N=7-9 replicates from 3 subjects. Each subject contained up to 3 technical replicates. Asterisk indicates p<0.05 by two-way ANOVA.

## Discussion

Burn wound conversion influences the healing potential of burn wounds and is associated with secondary morbidities such as wound infection.^[6]^ However, the mechanistic basis for burn wound conversion and its precise link to wound infection remain unclear. Here we identify that keratinocyte behavior promotes burn wound conversion following injury and is associated with wound infection. We find that microbial colonization of viable zone 2 tissue, analogous to the zone of stasis in humans, is significantly greater in larvae that exhibit increased epithelial damage resulting from keratinocyte migration. In contrast, preventing keratinocyte migration by treatment with isotonic medium results in significantly reduced colonization of viable tissue, demonstrating the potential for isotonic medium treatment to limit burn wound infection.

Successful tissue repair requires the coordinated migration of keratinocytes^[39, 40]^, and we previously found that burn injury induces dysregulated keratinocyte migration compared to mechanical tissue damage in zebrafish^[19]^. Unlike the migratory response to mechanical injury, in which keratinocytes are able to terminate migration within minutes upon reaching the wound edge, burn injury results in keratinocyte migration that lasts approximately 1 hour, suggesting the lack of a migratory stop signal. While our findings here link the process of keratinocyte migration to burn wound conversion, more work is needed to identify the mechanistic link between early keratinocyte behavior and widespread cell death that we observe in the tissue region analogous to the zone of stasis. An attractive hypothesis is that aberrant keratinocyte damage signaling promotes chronic inflammation, a process thought to contribute to burn wound conversion in human patients. Indeed, our laboratory has previously shown that burn wounding results in tissue-scale oxidative stress and chronic inflammation^[17, 19]^. Furthermore, we show here that isotonic medium treatment significantly reduces immune cell recruitment to burned tissue in sterile conditions, supporting a potential role for keratinocyte migration in mediating inflammatory signaling after tissue injury.

Epithelial damage and inflammation are also associated with wound infection in burn patients. Our data support this association in zebrafish, with robust *C. albicans* colonization evident in control larvae. Limiting epithelial damage by treatment with isotonic medium significantly reduced *C. albicans* colonization, with nearly 50% (12 out of 27) of treated larvae exhibiting no observable *C. albicans* by 24 hpb. These observations suggest that, at least in zebrafish, *C. albicans* colonization of burned tissue relies on viable keratinocytes physically migrating into zone 1 tissue where *C. albicans* initially exists. While we focused our current investigation *C. albicans*, our finding that isotonic medium treatment limits wound colonization by affecting the host-response – keratinocyte migration – not a fungal-specific pathway, leads us to hypothesize that isotonic medium may provide a general means of limiting burn wound infection across microbial flora.

While there are clear limitations in trying to apply our model of migration-mediated epithelial damage to human skin, our findings nonetheless demonstrate the potential for isotonic saline to improve burn wound healing. Aside from the obvious anatomical and environmental differences between larval zebrafish and human skin, there are distinct differences in the processes and timing of events that regulate tissue repair across species. For example, the lack of rapid keratinocyte migration in human skin, where keratinocytes migrate little and burn wounds heal comparatively slowly, highlights that a different mechanism must explain the observed benefit of topical saline treatment to burned human skin. Nevertheless, existing studies have explored the use of saline treatment in both mammalian models and burn patients. Treatment with saline showed promise when used as a washing agent in a small trial of pediatric burn patients^[41]^. One hour daily washing of the burned region with saline improved burn wound healing and reduced the likelihood of burn wound infection, albeit not to the level of statistical significance^[41]^. Despite the small sample of patients, this study showed the potential benefit of saline treatment and is generally consistent with our findings; however, based on our data it is tempting to speculate that saline treatment may have provided a greater benefit to patients if treatment was started earlier. While many patients were not seen for treatment until hours, if not days, following burning, our data suggest that the most impactful window for saline treatment would be immediately after wounding to abate progressive spread of damage in the zone of stasis. Highlighting the importance of the timing window for treatment with saline, a separate study showed saline was associated with mild patient discomfort.^[42]^ This trial involved long-term treatment in which patients were dressed with saline soaked bandages over several days. Importantly, this is not consistent with how our data suggests that saline treatment should be used to maximize the earliest benefits of limiting tissue damage.

In addition to determining the optimal timing for treatment of burn wounds, it is paramount to understand how potential treatment works at the molecular level. Our data suggests that isotonic solution may benefit burn wound healing by preserving the zone of stasis. Other studies have targeted burn wound conversion to improve burn wound healing, primarily by reducing inflammation and apoptotic cell death. One study comparing saline and platelet-rich plasma injected into the zone of stasis in burned rabbits found significantly less apoptosis when burns were injected with platelet-rich plasma. This resulted in a greater amount of vital tissue three days post-burn.^[43]^ Similarly, treatment with the phosphodiesterase inhibitor Udenafil improved burn wound healing in rats by limiting the spread of necrotic tissue and dampening inflammation into the zone of stasis.^[44]^ Interestingly, this study included saline controls; however, saline was administered orally, as opposed to topically as in our study. These studies highlight that a primary goal of burn wound therapeutics is to preserve the zone of stasis and that limiting cell death and inflammation is a common target. However, they also suggest potential species-specific differences in the wound healing process. For example, in zebrafish we found the contribution of apoptosis to cell death was relatively limited. Therefore, more work is needed to better understand how saline preserves the zone of stasis in zebrafish, prior to translating these findings to mammalian models.

The unique benefits of the zebrafish model organism lie in the ability to visualize the tissue response to burn injury at the cellular level, allowing new insights into burn wound pathology. Our future studies will extend this work to focus on the long-term impact of keratinocytes with the goal of identifying the conserved signaling pathways that link early migratory behavior with eventual death of cells in the zone of stasis. Identification of such pathways will benefit researchers working across the spectrum of model organisms. Finally, our work using zebrafish and cultured human skin suggests that immediate topical application of saline could be explored as a useful first aid intervention. An issue facing burn wound care providers is the lack of consensus on the best course of pre-hospitalization care for burn injuries.^[45–47]^ However, it is clear that a goal of such treatment should be to limit the spread of damage associated with burn wounds. Future studies aimed at better understanding how saline treatment prevents damage in mammalian skin, as well as optimizing the timing of saline treatment to limit burn wound damage are needed to achieve this goal.

## Acknowledgements

The authors would like to thank the University of Wisconsin-Madison, Department of Surgery Histology Resource Core (HRC), Dr. Angela Gibson, MD, PhD, FACS, PI of the HRC, along with certified Histotechnologists Sierra Raglin, HTL (ASCP)^CMP^ and Rebecca Lyons-Oelze, HTL (ASCP) for their assistance in the processing of frozen tissue samples. We also thank Dr. Rob Wheeler (University of Maine) for his gift of fluorescently labeled *Candida albicans* strains. Finally, we thank the members of the Huttenlocher and Gibson labs for their thoughtful input throughout the duration of this work. The authors acknowledge the following funding: K99 GM147303 to Adam Horn and R35 GM118027 to Anna Huttenlocher.

## Data Availability

Data will be made available upon request.

---

**Supplemental Figure 1: Isotonic medium does not prevent apoptotic cell death following burn injury.** (A) Images showing control and isotonic medium treated larvae after photoconversion of zone 1 tissue. Scale bar = 100 µm. (B) Quantification of zone 1 photoconverted area at the indicated time, hours post-burn (hpb). Data are normalized to photoconverted area 0 hpb. N = 10 larvae for each condition. (C) Images showing active-caspase-3 staining 6 hpb in control or isotonic medium treated larvae. Scale bar = 50 µm. (D) Quantification of active-Caspase-3 staining at the indicated time post-burn. N ≥ 19 control and 17 isotonic treated larvae for each time point. Asterisk indicates p<0.05 by two-way ANOVA.

**Supplemental Figure 2: *C. albicans* and host immune response to isotonic medium treatment.** (A) Representative image of mNeon-tagged *C. albicans* 24 hpb. Hyphal and yeast forms are present in burn wounded tissue. Scale bar = 10 µm. (B) *C. albicans* viability, measured by colony forming units (CFU) per milliliter. *C. albicans* was exposed to either control or isotonic water for 1 hour prior to viability plating. N = 3 independent experiments. (C) Images of *Tg(LyzCH2B-mCherryxMpeg1.1H2B-GFP)* larvae showing neutrophil (red) and macrophage (green) response to burn injury following either control or isotonic medium treatment. Solid white line delineates the wounded region, distal to the notochord, used for quantification. Scale bar = 100 µm. (D) Quantification of neutrophil recruitment at the indicated time post-burn. N ≥ 32 control and 34 isotonic treated larvae. (E) Quantification of macrophage recruitment at the indicated time post-burn. N ≥ 32 control and 34 isotonic treated larvae. Asterisk indicates p<0.05 by two-way ANOVA.

## References

1. Ogawa, R., Update on Hypertrophic Scar Management in Burn Patients. Clinics in Plastic Surgery, 2024. 51(3): p. 349–354.

2. Tsolakidis, S., et al., “Out of Touch”—recovering sensibility after burn injury: a review of the literature. European Burn Journal, 2022. 3(2): p. 370–376.

3. Gauglitz, G.G., et al., Hypertrophic scarring and keloids: pathomechanisms and current and emerging treatment strategies. Molecular medicine, 2011. 17: p. 113–125.

4. Salibian, A.A., et al., Current concepts on burn wound conversion—a review of recent advances in understanding the secondary progressions of burns. Burns, 2016. 42(5): p. 1025–1035.

5. Jackson, D.M., The diagnosis of the depth of burning. Journal of British Surgery, 1953. 40(164): p. 588–596.

6. Palackic, A., et al., Therapeutic strategies to reduce burn wound conversion. Medicina, 2022. 58(7): p. 922.

7. Kelly, E.J., et al., Infection and burn injury. European Burn Journal, 2022. 3(1): p. 165–179.

8. Wang, Y., et al., Burn injury: challenges and advances in burn wound healing, infection, pain and scarring. Advanced drug delivery reviews, 2018. 123: p. 3–17.

9. Williams, F.N., et al., The leading causes of death after burn injury in a single pediatric burn center. Critical care, 2009. 13: p. 1–7.

10. Gur, I., et al., Clinical impact of fungal colonization of burn wounds in patients hospitalized in the intensive care unit: a retrospective cohort study. Trauma Surgery & Acute Care Open, 2024. 9(1): p. e001325.

11. Capoor, M.R., et al., Fungal infections in burns: diagnosis and management. Indian Journal of Plastic Surgery, 2010. 43(S 01): p. S37–S42.

12. Jithendra, K., et al., Identification of fungal pathogens in burns patients with reference to Candida. Journal of Evidence based Medicine and Healthcare, 2015. 2: p. 1–8.

13. Sikwewa, K., et al., The occurrence of fungi from burn wound patients and antifungal susceptibility patterns: a cross-sectional study in Lusaka, Zambia. African Health Sciences, 2023. 23(3): p. 506–513.

14. Plichta, J.K., et al., Local burn injury impairs epithelial permeability and antimicrobial peptide barrier function in distal unburned skin. Critical care medicine, 2014. 42(6): p. e420–e431.

15. Martin, P. and R. Nunan, Cellular and molecular mechanisms of repair in acute and chronic wound healing. British Journal of Dermatology, 2015. 173(2): p. 370–378.

16. Le Guellec, D., G. Morvan-Dubois, and J.-Y. Sire, Skin development in bony fish with particular emphasis on collagen deposition in the dermis of the zebrafish (Danio rerio). International Journal of Developmental Biology, 2004. 48: p. 217–232.

17. Miskolci, V., et al., Distinct inflammatory and wound healing responses to complex caudal fin injuries of larval zebrafish. Elife, 2019. 8: p. e45976.

18. LeBert, D., et al., Damage-induced reactive oxygen species regulate vimentin and dynamic collagen-based projections to mediate wound repair. Elife, 2018. 7: p. e30703.

19. Fister, A.M., et al., Damage-induced basal epithelial cell migration modulates the spatial organization of redox signaling and sensory neuron regeneration. bioRxiv, 2024.

20. Houseright, R.A., et al., Myeloid-derived growth factor regulates neutrophil motility in interstitial tissue damage. Journal of Cell Biology, 2021. 220(8): p. e202103054.

21. Santos, D., S.M. Monteiro, and A. Luzio, General whole-mount immunohistochemistry of zebrafish (Danio rerio) embryos and larvae protocol. Teratogenicity Testing: Methods and Protocols, 2018: p. 365–371.

22. Bergeron, A.C., et al., Candida albicans and Pseudomonas aeruginosa interact to enhance virulence of mucosal infection in transparent zebrafish. Infection and Immunity, 2017. 85(11): p. 10.1128/iai.00475-17.

23. Gratacap, R.L., J.F. Rawls, and R.T. Wheeler, *Mucosal candidiasis elicits NF-*κ*B activation, proinflammatory gene expression and localized neutrophilia in zebrafish*. Disease models & mechanisms, 2013. 6(5): p. 1260–1270.

24. Wu, Y., et al., Microglia and amyloid precursor protein coordinate control of transient Candida cerebritis with memory deficits. Nature communications, 2019. 10(1): p. 58.

25. Liu, A., et al., Modeling early thermal injury using an ex vivo human skin model of contact burns. Burns, 2021. 47(3): p. 611–620.

26. Gibson, A.L. and S. Shatadal, A simple and improved method to determine cell viability in burn-injured tissue. Journal of Surgical Research, 2017. 215: p. 83–87.

27. Gault, W.J., B. Enyedi, and P. Niethammer, Osmotic surveillance mediates rapid wound closure through nucleotide release. Journal of Cell Biology, 2014. 207(6): p. 767–782.

28. Sforna, L., et al., *Piezo1 controls cell volume and migration by modulating swelling*-*activated chloride current through Ca2+ influx*. Journal of Cellular Physiology, 2022. 237(3): p. 1857–1870.

29. Gurskaya, N.G., et al., Engineering of a monomeric green-to-red photoactivatable fluorescent protein induced by blue light. Nature biotechnology, 2006. 24(4): p. 461–465.

30. Singer, A.J., et al., Apoptosis and necrosis in the ischemic zone adjacent to third degree burns. Academic Emergency Medicine, 2008. 15(6): p. 549–554.

31. Rennekampff, H.-O. and Z. Alharbi, Burn injury: mechanisms of keratinocyte cell death. Medical Sciences, 2021. 9(3): p. 51.

32. Schofield, C.M., et al., Correlation of culture with histopathology in fungal burn wound colonization and infection. Burns, 2007. 33(3): p. 341–346.

33. Desai, J.V., Candida albicans hyphae: from growth initiation to invasion. Journal of Fungi, 2018. 4(1): p. 10.

34. O’sullivan, S. and T. O’connor, Immunosuppression following thermal injury: the pathogenesis of immunodysfunction. British journal of plastic surgery, 1997. 50(8): p. 615–623.

35. Church, D., et al., Burn wound infections. Clinical microbiology reviews, 2006. 19(2): p. 403–434.

36. Jeschke, M.G., et al., Burn injury. Nature reviews Disease primers, 2020. 6(1): p. 11.

37. Kennard, A.S. and J.A. Theriot, Osmolarity-independent electrical cues guide rapid response to injury in zebrafish epidermis. Elife, 2020. 9: p. e62386.

38. Gillenwater, J. and W. Garner, Acute fluid management of large burns: pathophysiology, monitoring, and resuscitation. Clinics in plastic surgery, 2017. 44(3): p. 495–503.

39. Peña, O.A. and P. Martin, Cellular and molecular mechanisms of skin wound healing. Nature Reviews Molecular Cell Biology, 2024: p. 1–18.

40. Rousselle, P., F. Braye, and G. Dayan, Re-epithelialization of adult skin wounds: Cellular mechanisms and therapeutic strategies. Advanced Drug Delivery Reviews, 2019. 146: p. 344–365.

41. Oyelami, O., et al., Management of Burn Injuries by Daily soaking in Normal Saline prior to Dressing. Nigerian Journal of Paediatrics, 2001. 28(4): p. 115–118.

42. Hindy, A., Comparative study between sodium carboxymethyl-cellulose silver, moist exposed burn ointment, and saline-soaked dressing for treatment of facial burns. Annals of Burns and Fire Disasters, 2009. 22(3): p. 131.

43. Uraloğlu, M., et al., The effect of platelet-rich plasma on the zone of stasis and apoptosis in an experimental burn model. Plastic Surgery, 2019. 27(2): p. 173–181.

44. Ural, A., et al., The Effect of Udenafil on Stasis Zone in an Experimental Burn Model. Annals of Plastic Surgery, 2022. 88(1): p. 38–43.

45. Goodwin, N.S., Just the tip of the iceberg-Inconsistent information on a global scale and the need for a" standard" model of burn 1st aid. Burns: Journal of the International Society for Burn Injuries, 2019. 45(3): p. 746–748.

46. Goodwin, N.S., *Burn first aid issues again—“Not seeing the forest for the trees”*. burns, 2021. 47(4): p. 970.

47. Cuttle, L., et al., A review of first aid treatments for burn injuries. Burns, 2009. 35(6): p. 768–775.

