## Supplementary figures and images for "Isotonic medium treatment limits burn wound microbial colonization and improves tissue repair"

### Supplemental Figure 1

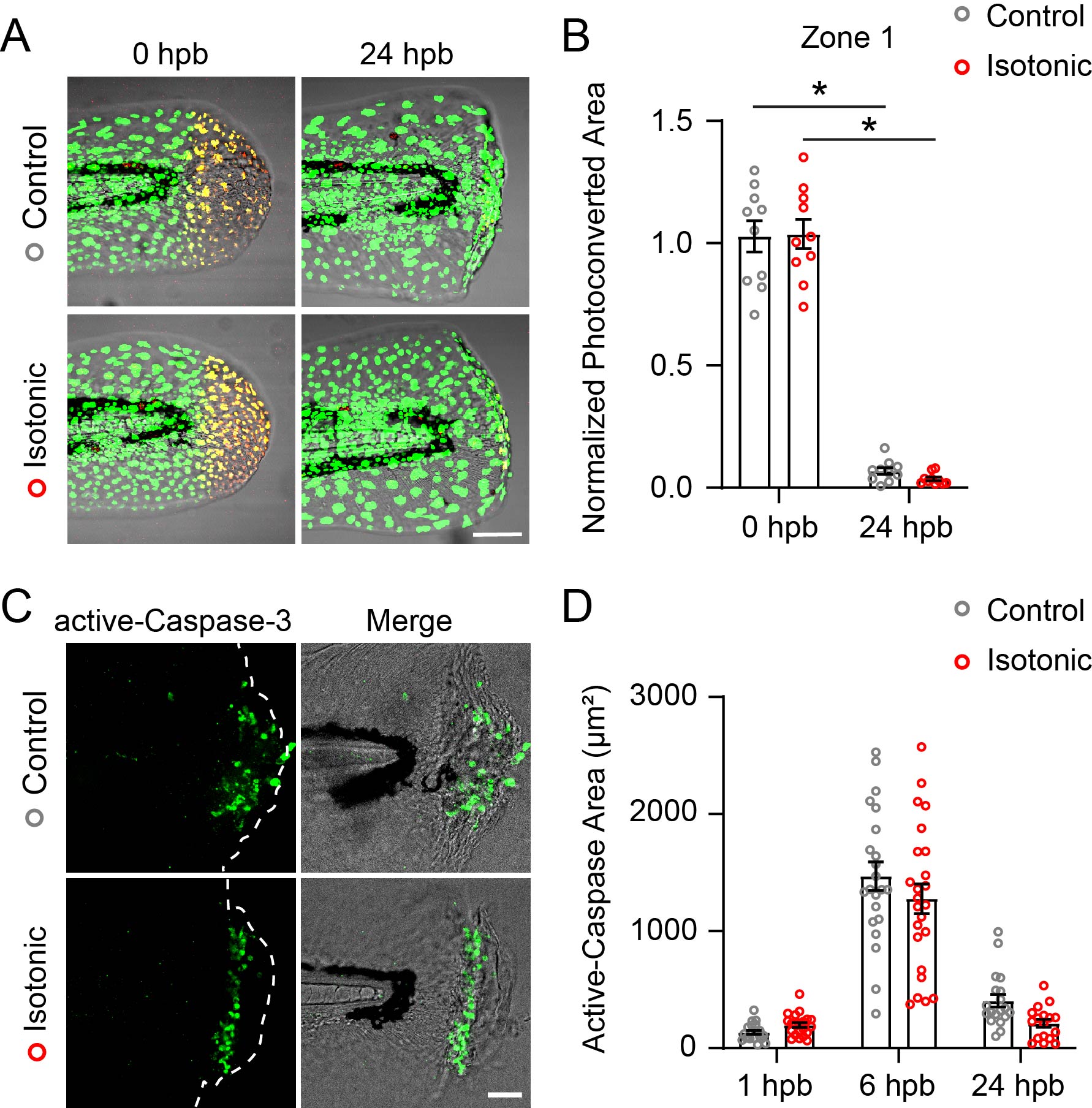

### Supplemental Figure 2

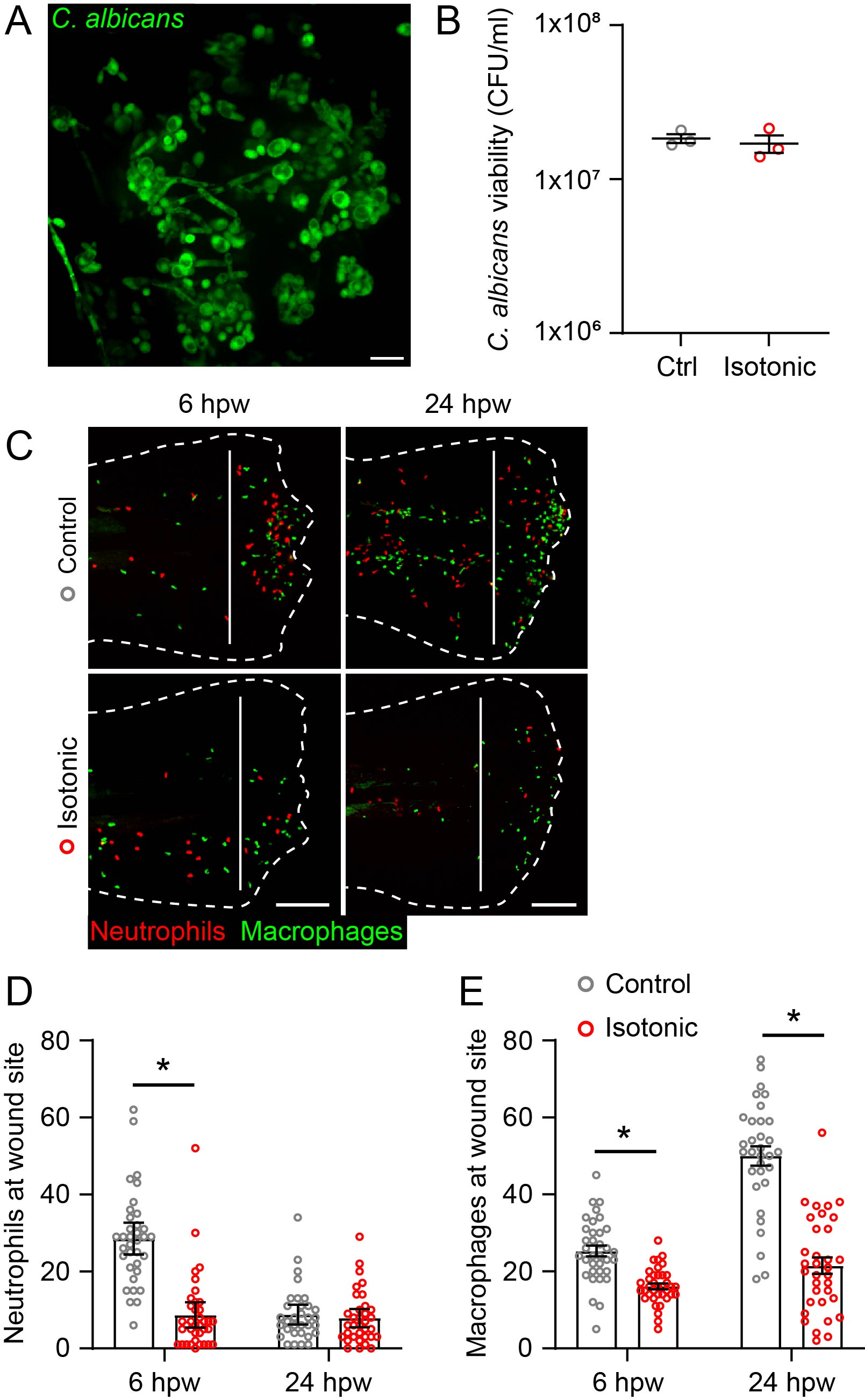
